# Water and temperature stress define the optimal flowering period for wheat in south-eastern Australia

**DOI:** 10.1101/115287

**Authors:** B.M. Flohr, J.R. Hunt, J.A. Kirkegaard, J.R. Evans

## Abstract

Across the Australian wheat belt, the time at which wheat flowers is a critical determinant of yield. In all environments an optimal flowering period (OFP) exists which is defined by decreasing frost risk, and increasing water and heat stress. Despite their critical importance, OFPs have not been comprehensively defined across south eastern Australia’s (SEA) cropping zone using yield estimates incorporating temperature, radiation and water-stress. In this study, the widely validated cropping systems model APSIM was used to simulate wheat yield and flowering date, with reductions in yield applied for frost and heat damage based on air temperatures during sensitive periods. Simulated crops were sown at weekly intervals from April 1 to July 15 of each year. The relationship between flowering date and grain yield was established for 28 locations using 51-years (1963-2013) of climate records. We defined OFPs as the flowering period which was associated with a mean yield of ≥ 95% of maximum yield from the combination of 51 seasons and 16 sowing dates. OFPs for wheat in SEA varied with site and season and were largely driven by seasonal water supply and demand, with extremes of heat and temperature having a secondary though auto-correlated effect. Quantifying OFPs will be a vital first step to identify suitable genotype x sowing date combinations to maximise yield in different locations, particularly given recent and predicted regional climate shifts including the decline in autumn rainfall.

## 1. Introduction

In all environments there exists a period during which wheat (*Triticum aestivum* L.) must flower in order for grain yield to be maximised, herein referred to as the optimal flowering period (OFP). Flowering during the optimal period is critical to grain yield as grain number is determined just prior to and at flowering (Fischer, 1985) and grain yield is most sensitive to stresses during this period, including drought (del Moral *et al.*, 2003; Giunta *et al.*, 1993) and extreme high (Ferris *et al.*, 1998; Shpiler and Blum, 1990; Tashiro and Wardlaw, 1990) and low temperatures (Boer *et al.*, 1993; Fuller *et al.*, 2007). In temperate climates such as northern Europe, flowering date has a broad optimum. However, in environments with a distinct dry season, flowering outside narrow OFPs can result in drastic yield reductions (Bodner *et al.*, 2015). The wheat belt of south eastern Australia (SEA) is one such environment, which has a predominantly Mediterranean climate with a cool wet season during which rain-fed wheat and other grain crops are grown, and a hot, dry season where land is left fallow. Whilst rainfall in the north-east of the region is equi-seasonal in distribution, cropping is still confined to the cool season by high summer temperatures and insufficient precipitation to sustain summer crops (Chenu *et al.*, 2013; Potgieter *et al.*, 2002). In the 2012/ 2013 season the south eastern states of Australia (New South Wales, Victoria and South Australia) produced over 14 Mt of wheat, 63% of Australia’s total wheat production (Commonwealth of Australia, 2013). The majority of annual production is exported, making the region important for global food security.

In SEA, spring wheat cultivars are established following rainfall in April-May (austral autumn) and grow during winter to mature at the end of spring. Significant yield progress has been made by breeders selecting cultivars with development patterns such that once established in autumn they will flower during the optimal period (Richards, 1991; Richards *et al.*, 2014). However, since 1996, rains that could once be relied upon by farmers to establish crops in April-May have declined significantly (Cai *et al.*, 2012; Pook *et al.*, 2009). This decline was particularly severe during the millennium drought (Verdon-Kidd *et al.*, 2014) at which time wheat crops established and flowered too late and yield was reduced by terminal drought and heat (Commonwealth of Australia, 2013). Reduced autumn rainfall has been attributed to anthropogenic climate change (Cai *et al.*, 2012; Murphy and Timbal, 2008) and is likely to persist. New combinations of management and genetics will be required to stabilise flowering date in order to overcome the observed yield decline (Kirkegaard and Hunt, 2010), and maintain the viability of SEA wheat farms and their contribution to global food security. A clear first step in this process is to identify the current OFP for environments in the SEA wheat belt.

A combination of environmental factors (precipitation, soil type, temperature) influence the opening, closing and duration of the OFP. Previous authors have stated that OFPs in SEA occur after the last spring frost and before the onset of heat and water stress (Anderson and Smith, 1990; Richards, 1991; Zheng *et al.*, 2012). Untimely spring frosts (September to October) are common in the Australian wheat belt (Boer *et al.*, 1993; Fuller *et al.*, 2007; Zheng *et al.*, 2012). A yield penalty of 10 % as a direct result of frost is common (Fuller *et al.*, 2007), and more catastrophic events are frequent (Crimp *et al.*, 2015). Zheng *et al.* (2015) analysed the frost and heat patterns of the Australian wheat belt, and found that the only regions that could be classified as almost “frost free” were some areas of the coastline in South Australia and north-east of central Queensland, while frosts occurred in other regions in 80% of years. Wheat is most sensitive to frost during reproductive growth stages. When wheat ears are exposed to freezing temperature after heading, frost damage will reduce the number of grain and sometimes cause death of entire ears (Fuller *et al.*, 2007).

High temperatures during sensitive reproductive growth stages can also result in a yield penalty (Ferris *et al.*, 1998). Gomez-Macpherson and Richards (1995) found that grain yield declined by 1.3% per day that sowing was delayed after late-May due to high temperatures around the time of anthesis and grain-fill. High temperature events (>35^o^C) during the period between head emergence to 10 days after anthesis can significantly reduce grain number and quality (Tashiro & Wardlaw, 1990). Similarly, heat shock during the grain filling period can also cause grain abortion and degrade grain quality (Randall and Moss, 1990; Stone and Nicolas, 1995).

Perhaps the most important determinant of the OFP is the pattern of water supply and demand experienced in a given environment (Bodner *et al.*, 2015). Whilst drought patterns in SEA are well described (Chenu *et al.*, 2013), the effect of seasonal water supply and demand in determining OFPs has been overlooked in previous analyses of OFPs e.g. Zheng *et al.* (2012), Zheng *et al.* (2013), Bell *et al.* (2015). In all of these studies OFPs were defined only by temperature extremes, which ignores the critical role that water supply and demand plays in defining OFPs in SEA.

To obtain an accurate definition of the OFP for a specific environment, Anderson and Smith (1990) suggest that time of sowing experiments should be conducted over a range of seasons. Anderson *et al.* (1996) defined the optimal flowering period from field experiments by “…using the mean flowering dates …of the sites and the optimum flowering period was taken to be 10 days either side of the mean optimum date [for grain yield]”. Experiments like these are expensive, and the recommendations to farmers are specific to the experimental conditions (e.g. temperature, rainfall and soil water holding capacity) during the period of the experiments, and may not reflect long-term climatic patterns (Asseng *et al.*, 2001). More recently, Zheng (2012) analysed heat and frost patterns of the wheat belt to calculate flowering windows based on occurrence of last frost days and first heat days.

Alternatively, an analysis of historic climate records using a crop simulator such as APSIM (Holzworth *et al.*, 2014; Keating *et al.*, 2003) allows one to identify management strategies to achieve optimum sowing date and flowering period (Asseng *et al.*, 2001; Zheng *et al.*, 2012) and can account for both temperature and water stress simultaneously. This is especially useful in the seasonally variable production environments of SEA (Asseng and Pannell, 2013; Asseng *et al.*, 2001; Turner, 2004).

This study sought to define OFPs in SEA to assess management and genetic interventions to overcome yield reductions due to the decline in autumn rainfall. It uses APSIM to incorporate the seasonal effects of water supply and demand, radiation and temperature on grain yield. By using the potential yield predictions to integrate the effects of temperature and radiation as well as water supply and demand, it extends the work of Zheng *et al.* (2012) who used only air temperature records alone (i.e. frost, heat) to define OFPs for the entire Australian wheat belt.

## 2. Materials and Methods

### 2.1 Site selection and crop simulation approach

Locations were selected to represent environments where wheat is grown in the cropping belt of SEA (Fig. 1, Table 1), and based on the availability of accurate soil characterization from the APSoil database (Dalgliesh *et al.*, 2009) and patched-point meteorological weather stations from the SILO database (Jeffrey *et al.*, 2001). At some sites (Hopetoun, Swan Hill and Bogan Gate), two different soil files were selected to compare the effect of soil type on the OFP. The cropping systems model Agricultural Production Systems SIMulator (APSIM), version 7.6 (Holzworth *et al.*, 2014; Keating *et al.*, 2003) was used to simulate wheat flowering date and yield using 51 years (1963- 2013) of climate data. Simulation of wheat growth, development and yield in APSIM has been extensively validated in numerous studies across southern Australia (Asseng *et al.*, 2001; Carberry *et al.*, 2009; Hochman *et al.*, 2009; Lilley *et al.*, 2003; Lilley and Kirkegaard, 2007), and no further validation was undertaken here. The key APSIM modules used in the analysis were Wheat (wheat crop growth and development) and Manager (specifying sowing rules).

**Figure 1:**
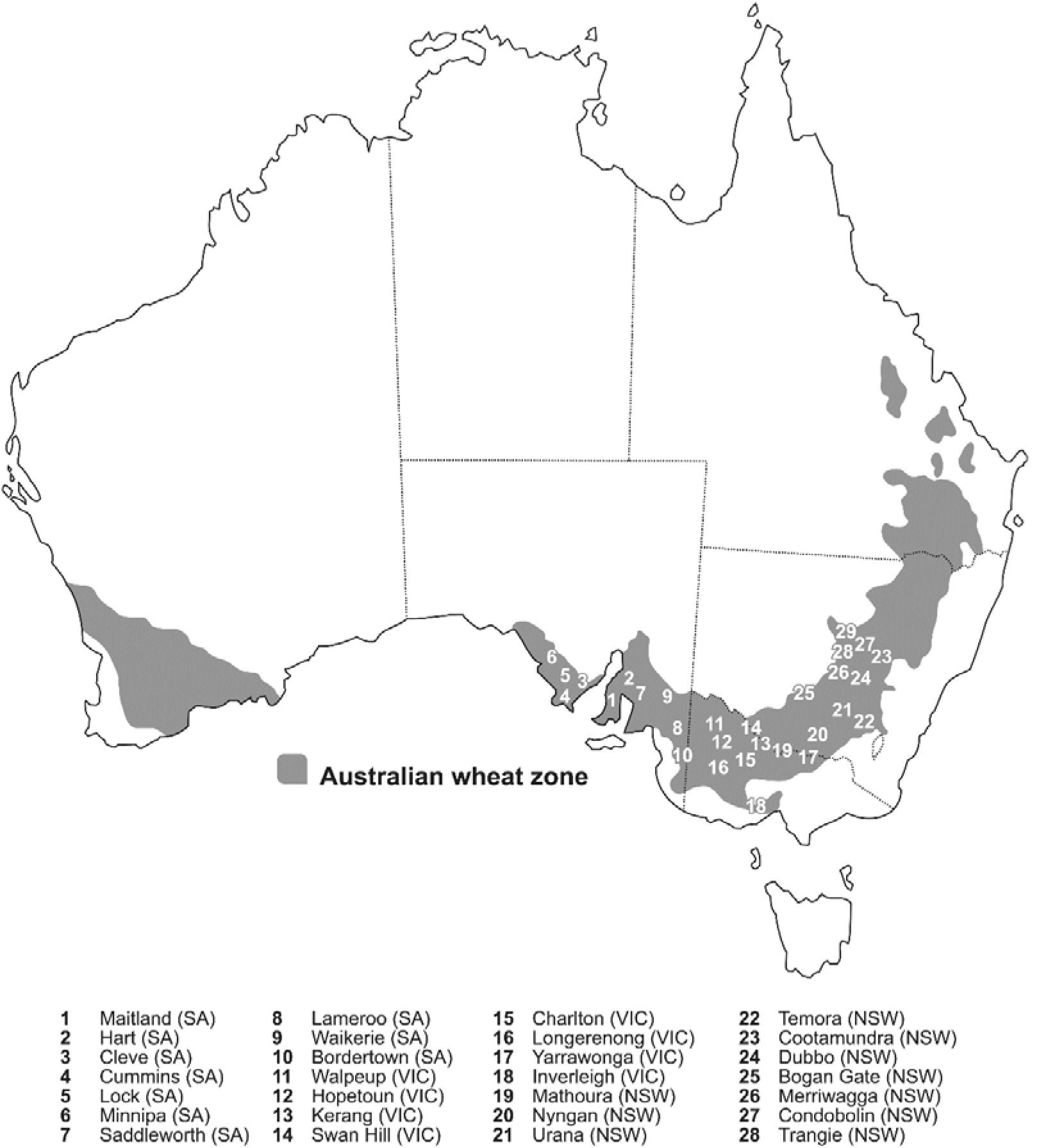
Dry-land cereal-cropping areas in Australia (shaded) and the locations used for this study.

**Table 1:**
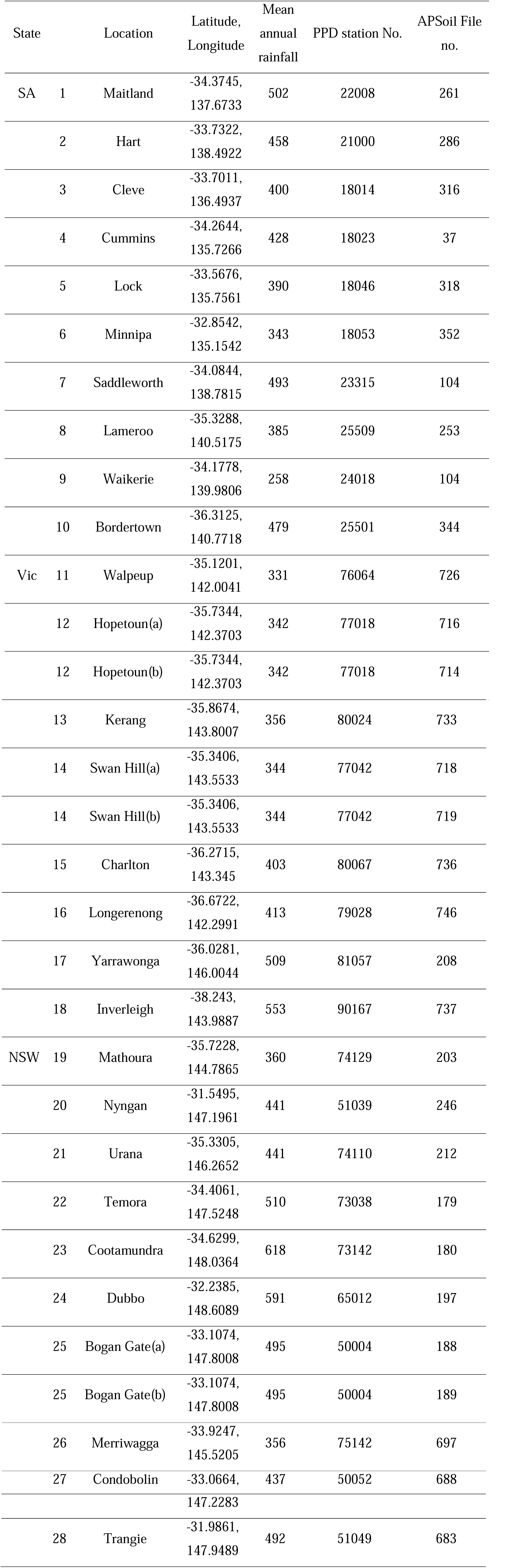
The 28 locations, where (a) and (b) refer to different soil types at the same location, used in the study and corresponding long-term mean annual rainfall, patch point dataset (PPD) station number, APSoil file number used in the simulation study.

| State |  | Location | Latitude,<br>Longitude | Mean<br>annual<br>rainfall | PPD station No. | APSoil File<br>no. |
| --- | --- | --- | --- | --- | --- | --- |
| SA | 1 | Maitland | -34.3745,<br>137.6733 | 502 | 22008 | 261 |
|  | 2 | Hart | -33.7322,<br>138.4922 | 458 | 21000 | 286 |
|  | 3 | Cleve | -33.7011,<br>136.4937 | 400 | 18014 | 316 |
|  | 4 | Cummins | -34.2644,<br>135.7266 | 428 | 18023 | 37 |
|  | 5 | Lock | -33.5676,<br>135.7561 | 390 | 18046 | 318 |
|  | 6 | Minnipa | -32.8542,<br>135.1542 | 343 | 18053 | 352 |
|  | 7 | Saddleworth | -34.0844,<br>138.7815 | 493 | 23315 | 104 |
|  | 8 | Lameroo | -35.3288,<br>140.5175 | 385 | 25509 | 253 |
|  | 9 | Waikerie | -34.1778,<br>139.9806 | 258 | 24018 | 104 |
|  | 10 | Bordertown | -36.3125,<br>140.7718 | 479 | 25501 | 344 |
| Vic | 11 | Walpeup | -35.1201,<br>142.0041 | 331 | 76064 | 726 |
|  | 12 | Hopetoun(a) | -35.7344,<br>142.3703 | 342 | 77018 | 716 |
|  | 12 | Hopetoun(b) | -35.7344,<br>142.3703 | 342 | 77018 | 714 |
|  | 13 | Kerang | -35.8674,<br>143.8007 | 356 | 80024 | 733 |
|  | 14 | Swan Hill(a) | -35.3406,<br>143.5533 | 344 | 77042 | 718 |
|  | 14 | Swan Hill(b) | -35.3406,<br>143.5533 | 344 | 77042 | 719 |
|  | 15 | Charlton | -36.2715,<br>143.345 | 403 | 80067 | 736 |
|  | 16 | Longerenong | -36.6722,<br>142.2991 | 413 | 79028 | 746 |
|  | 17 | Yarrawonga | -36.0281,<br>146.0044 | 509 | 81057 | 208 |
|  | 18 | Inverleigh | -38.243,<br>143.9887 | 553 | 90167 | 737 |
| NSW | 19 | Mathoura | -35.7228,<br>144.7865 | 360 | 74129 | 203 |
|  | 20 | Nyngan | -31.5495,<br>147.1961 | 441 | 51039 | 246 |
|  | 21 | Urana | -35.3305,<br>146.2652 | 441 | 74110 | 212 |
|  | 22 | Temora | -34.4061,<br>147.5248 | 510 | 73038 | 179 |
|  | 23 | Cootamundra | -34.6299,<br>148.0364 | 618 | 73142 | 180 |
|  | 24 | Dubbo | -32.2385,<br>148.6089 | 591 | 65012 | 197 |
|  | 25 | Bogan Gate(a) | -33.1074,<br>147.8008 | 495 | 50004 | 188 |
|  | 25 | Bogan Gate(b) | -33.1074,<br>147.8008 | 495 | 50004 | 189 |
|  | 26 | Merriwagga | -33.9247,<br>145.5205 | 356 | 75142 | 697 |
|  | 27 | Condobolin | -33.0664, | 437 | 50052 | 688 |

|  |  |  |  |  |  |
| --- | --- | --- | --- | --- | --- |
|  |  | 147.2283 |  |  |  |
| 28 | Trangie | -31.9861,<br>147.9489 | 492 | 51049 | 683 |

APSIM calculates flowering date i.e. anthesis (and other crop stages) by accumulation of thermal time. In the APSIM plant module, thermal time is calculated from the average of 3-hourly air temperatures estimated using diurnal cycling between recorded maximum and minimum temperatures, with the eight 3- hour estimates averaged to determine the daily value of thermal time for the day. The length of each crop stage between emergence and floral initiation is determined by the accumulation of thermal time, and cultivar- specific factors accounting for vernalisation and photoperiod responses (Ritchie and NeSmith, 1991). The length of each crop stage from floral initiation to maturity is determined only by accumulation of thermal time. The root-mean-square error (RMSE) in the ability for APSIM-Wheat to predict flowering time is 6.2 days (Zheng *et al.*, 2013).

### 2.2 Crop management set up

All simulated crops were sown at 150 plants/m^2^, at a depth of 30 mm with a row spacing of 300 mm. In APSIM cultivars are allocated a vernalisation and photoperiod “factor” which represent sensitivity to environment elements cold and day length where high values are more sensitive. The model then uses these factors to calculate cultivar specific developmental rates. The cultivar parameters selected here represent a spring wheat of mid to fast phenological development, typical of varieties grown in SEA (e.g. Scout, Spitfire, Mace etc.). This was based on the APSIM base cultivar with vernalisation sensitivity of 1.5 and photoperiod sensitivity of 3.0. For comparison, a winter wheat (e.g. Wedgetail) with vernalisation sensitivity of 4, photoperiod sensitivity of 0, and an intermediate facultative wheat (e.g. Eaglehawk) with a vernalisation sensitivity of 2.5 and photoperiod sensitivity of 4 were also simulated at selected sites (Temora, NSW and Lameroo, S.A) under the same management parameters to observe any change in yield and OFP.

APSIM-Manager was used to sow a crop on a fixed date at weekly intervals from 1 April to 15 July of each year. Nitrogen was applied as NO_3_ with a fertilizer rule, which was maintained above 100 kg/ha in the top three layers of the soil throughout the season such that nitrogen supply did not limit yield. In the simulation, the initial plant available water was set to 0 mm, and the crop received 15 mm of irrigation at sowing to ensure that it would emerge shortly after it was sown. APSIM assumes crops are grown free of weeds and disease. As highlighted in the recent review by Barlow *et al.* (2015), APSIM does not currently account for frost and heat events in its yield predictions. Therefore, a reduction for frost and heat damage based on air temperature obtained from patched point meteorological weather stations was applied as per Bell *et al.* (2015). To estimate the effect of heat or frost stress events on grain yield during sensitive growth stages, temperature ranges were categorized into mild, medium and severe stress with a corresponding impact on yield during different growth stages, these estimations are based on the literature and expert opinion (Bell *et al.*, 2015) (Table 2). Yield reductions were cumulative for multiple events that occurred during the sensitive stages of plant growth (Table 2). The combination of management rules and frost and heat rules ensure that the OFP was defined for each environment by the combination the drought pattern, temperature and radiation. Outputs from the simulation were annual potential grain yield and annual grain yield modified for frost and heat damage, referred to as the frost and heat limited (FHL) yield hereafter, at different sowing dates and corresponding flowering dates. To quantify the effect of stress on the OFP, the mean stress index of frost and heat, and the mean water stress between floral initiation and maturity applied to the crop was plotted for selected sites, with 0 indicating zero effect on grain yield, and 1 having the greatest effect on grain yield.

**Table 2:** Temperature criteria for frost and heat stress during sensitive Zadoks growth stages and corresponding estimated yield reduction, from Bell *et al.* (2015). Yield reduction was calculated for each day and multiplied, so that increasing numbers of stress events results in cumulative reduction in yield.

| Temperature | Stress level | Zadoks sensitive stage | Yield reduction per day |
| --- | --- | --- | --- |
| 0 to 2<sup>0</sup>C | Mild | 60-69 | 10% |
| -2 to 0<sup>0</sup>C | Moderate | 60-75 | 20% |
| < -2<sup>0</sup>C | Severe | 60-79 | 90% |
| 32 to 34<sup>0</sup>C | Mild | 60-79 | 10% |
| 34 to 36<sup>0</sup>C | Moderate | 60-79 | 20% |
| >36<sup>0</sup>C | Severe | 60-79 | 30% |

For each location, over the 51 years x 16 sowing dates of simulation, the FHL yield was attributed to the predicted flowering date in each simulation. OFPs were defined by calculating a 15-day running mean of FHL yield over the 51 years of simulation. Flowering dates corresponding to ≥ 95% of the peak mean FHL yield defined the dates of the OFP for each location. Three lines were plotted, the 15-day running mean, the positive standard deviation, and the negative standard deviation associated with the running mean. To demonstrate how the OFP changes from season to season, the 15-day running mean of yield was split into 10, 20, 30, 40, 50, 60, 70, 80 and 90^th^ percentiles for selected locations.

The APSIM simulation was also used to estimate the optimal sowing date range for each location to achieve the OFPs. To define the sowing range, the sowing dates that corresponded to the flowering dates that achieved the highest 15-day running mean yield were split into earliest sowing date, 25^th^ percentile, median, 75^th^ percentile and latest sowing dates.

To observe effects of recent rainfall decline and increasing temperatures within the 51-year simulation, APSIM output for yield and flowering date was analysed for two time periods 1963-1997 and 1998-2013. The 15-year time period (1998-2013) includes the period of April-May rainfall decline experienced at many of the sites in the south-eastern wheat belt described by Cai *et al.* (2012).

## 3. Results

### 3.1 Defining optimal flowering periods

Yield and OFP varied across 28 locations according to the temperature, radiation and rainfall patterns and soil type of each environment (Table 3). The highest peak mean FHL yield was in Maitland (4910 kg ha^-1^), followed by Inverleigh (4841 kg ha^-1^). The lowest yielding location was Waikerie (1819 kg ha^-1^). The open and close dates and duration varied significantly across the wheat belt. The earliest OFP open date was at Minnipa (22 August), while the latest was at Inverleigh (12 October). The longest and shortest durations of the OFP were Inverleigh (26 days) and Nyngan (4 days), and the latest and earliest close dates were Inverleigh (6 November) and Nyngan and Waikerie (29 August). Timing of the OFP was related to mean annual rainfall (Fig. 2) and consequently yield. For example, Inverleigh had a mean annual rainfall of 553 mm, and an OFP from 12 October to 6 November. In contrast, Waikerie had a mean annual rainfall of 258 mm and an earlier OFP from 23 August to the 29 August. However, high annual rainfall often coincided with a cooler growing season in SEA (data not shown) e.g. Inverleigh, so effects of temperature and water availability are confounded.

**Table 3:** Optimal flowering periods (OFPs), peak mean of frost and heat limited yield (kg ha^-1^) and corresponding flowering date and sowing date range for a mid-fast cultivar over 51-years (1963-2013), for the 28 locations where (a) and (b) refer to different soil types at the same location, ranked in alphabetical order.

| Location | Optimal flowering period |  | Peak mean yield (kg ha<sup>-1</sup>) and corresponding flowering date |  | Sowing date range for corresponding peak mean |  |  |  |  |
| --- | --- | --- | --- | --- | --- | --- | --- | --- | --- |
|  | Open | Close |  |  | Earliest | 25<sup>th</sup> percentile | Median | 75<sup>th</sup> percentile | Latest |
| Bogan Gate (a) | 19-Sep | 3-Oct | 4283 | 27-Sep | 12-May | 16-May | 21-May | 27-May | 1-Jun |
| Bogan Gate (b) | 18-Sep | 1-Oct | 3650 | 21-Sep | 6-May | 8-May | 13-May | 17-May | 21-May |
| Bordertown | 26-Sep | 15-Oct | 3577 | 7-Oct | 16-May | 23-May | 29-May | 3-Jun | 9-Jun |
| Charlton | 21-Sep | 30-Sep | 2920 | 23-Sep | 29-Apr | 6-May | 7-May | 10-May | 14-May |
| Cleve | 7-Sep | 18-Sep | 3402 | 13-Sep | 13-May | 15-May | 20-May | 24-May | 30-May |
| Condobolin | 11-Sep | 19-Sep | 2435 | 15-Sep | 29-Apr | 2-May | 7-May | 10-May | 15-May |
| Cootamundra | 6-Oct | 20-Oct | 4338 | 12-Oct | 6-May | 13-May | 20-May | 24-May | 31-May |
| Cummins | 8-Sep | 28-Sep | 3843 | 18-Sep | 15-May | 19-May | 24-May | 27-May | 1-Jun |
| Dubbo | 15-Sep | 22-Sep | 3881 | 18-Sep | 6-May | 7-May | 11-May | 15-May | 18-May |
| Hart | 21-Sep | 2-Oct | 4242 | 24-Sep | 6-May | 8-May | 13-May | 18-May | 22-May |
| Hopetoun (a) | 7-Sep | 11-Sep | 2900 | 8-Sep | 1-May | 5-May | 10-May | 14-May | 19-May |
| Hopetoun (b) | 4-Sep | 9-Sep | 1903 | 5-Sep | 20-Apr | 22-Apr | 25-Apr | 29-Apr | 30-Apr |
| Inverleigh | 12-Oct | 6-Nov | 4841 | 22-Oct | 2-Jun | 11-Jun | 15-Jun | 24-Jun | 1-Jul |
| Kerang | 12-Sep | 18-Sep | 3131 | 15-Sep | 25-Apr | 29-Apr | 3-May | 5-May | 9-May |
| Lameroo | 28-Aug | 20-Sep | 3154 | 7-Sep | 26-Apr | 29-Apr | 2-May | 6-May | 11-May |
| Lock | 4-Sep | 14-Sep | 3049 | 9-Sep | 6-May | 7-May | 13-May | 15-May | 20-May |
| Longerenong | 6-Oct | 10-Oct | 2749 | 7-Oct | 9-May | 12-May | 16-May | 25-May | 27-May |
| Maitland | 16-Sep | 9-Oct | 4910 | 1-Oct | 27-May | 29-May | 5-Jun | 10-Jun | 14-Jun |
| Mathoura | 15-Sep | 22-Sep | 2268 | 18-Sep | 26-Apr | 30-Apr | 3-May | 6-May | 9-May |
| Merriwagga | 27-Aug | 10-Sep | 2619 | 31-Aug | 22-Apr | 25-Apr | 27-Apr | 29-Apr | 6-May |
| Minnipa | 22-Aug | 8-Sep | 3514 | 25-Aug | 29-Apr | 30-Apr | 6-May | 6-May | 8-May |
| Nyngan | 26-Aug | 29-Aug | 2234 | 27-Aug | 27-Apr | 30-Apr | 2-May | 7-May | 9-May |
| Saddleworth | 9-Sep | 24-Sep | 3857 | 17-Sep | 6-May | 10-May | 15-May | 17-May | 23-May |
| Swan Hill (a) | 1-Sep | 20-Sep | 3673 | 15-Sep | 26-Apr | 2-May | 6-May | 9-May | 12-May |
| Swan Hill (b) | 1-Sep | 10-Sep | 2805 | 5-Sep | 19-Apr | 24-Apr | 27-Apr | 30-Apr | 3-May |
| Temora | 25-Sep | 10-Oct | 3038 | 3-Oct | 4-May | 6-May | 13-May | 18-May | 22-May |
| Trangie | 12-Sep | 30-Sep | 4746 | 22-Sep | 13-May | 14-May | 21-May | 25-May | 30-May |
| Urana | 18-Sep | 29-Sep | 3282 | 23-Sep | 30-Apr | 4-May | 8-May | 12-May | 16-May |
| Waikerie | 23-Aug | 29-Aug | 1819 | 25-Aug | 22-Apr | 23-Apr | 27-Apr | 29-Apr | 3-May |
| Walpeup | 8-Sep | 17-Sep | 3318 | 11-Sep | 29-Apr | 1-May | 4-May | 10-May | 13-May |
| Yarrawonga | 25-Sep | 2-Oct | 3594 | 28-Sep | 30-Apr | 2-May | 8-May | 13-May | 17-May |

**Figure 2:**
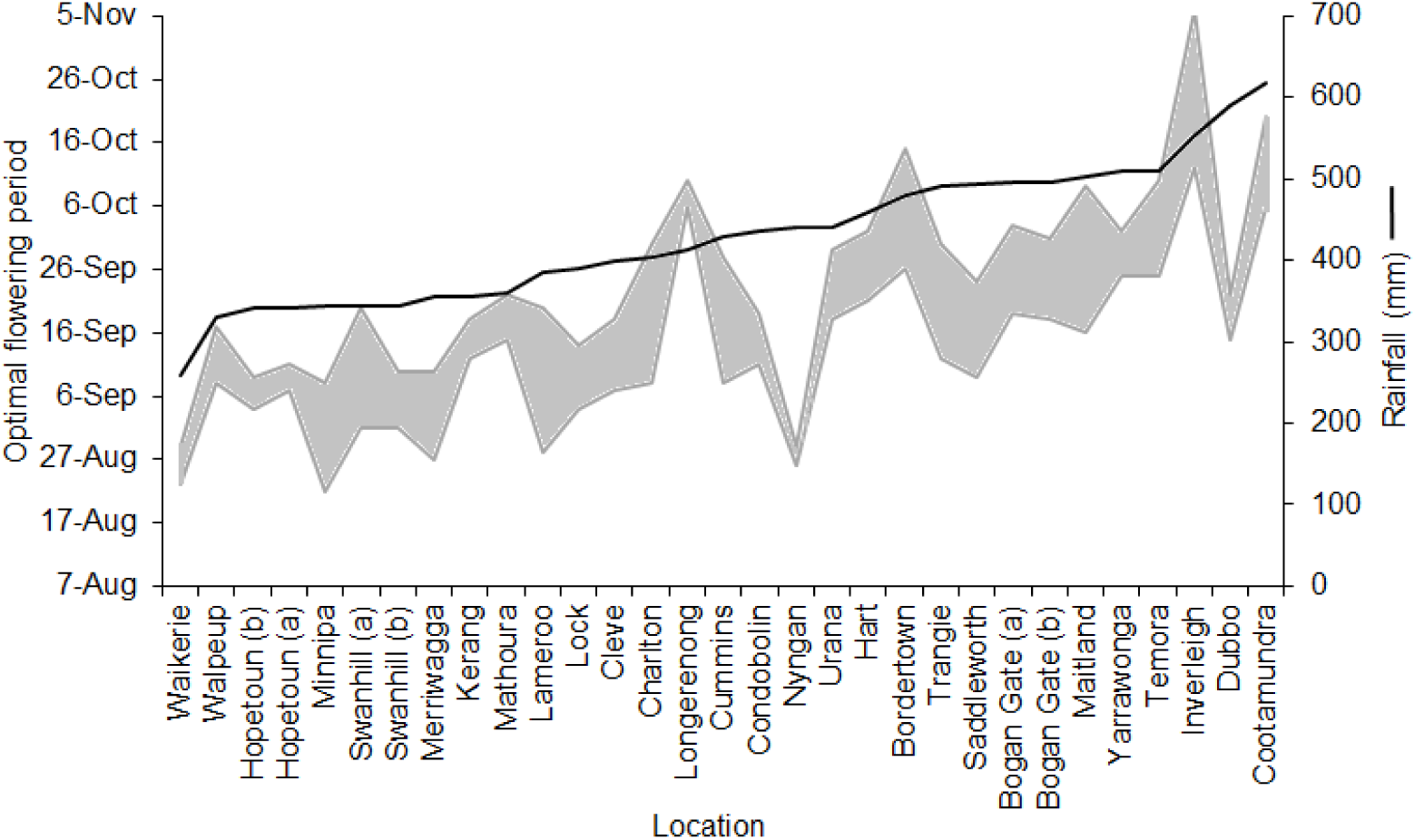
The relationship between optimal flowering period (shaded grey) of a mid-fast cultivar of wheat (Table 3), and mean annual rainfall (mm) (line) (Table 2) for 28 locations, where (a) and (b) refer to different soil types at the same location.

10^th^- 90^th^ percentiles of FHL yield simulated at each flowering date (Fig. 3) reveal the seasonal variability in the OFP. Depending on which environment factor had the greatest effect on yield at each location i.e. frost, radiation, heat or drought, the OFP shifted accordingly. At some locations, e.g. Lameroo (Fig. 3A), in low yielding seasons the OFP was earlier than in more favourable seasons, as early flowering allowed crops to escape spring drought. At other locations with high incidence of frost, e.g. Temora (Fig. 3B), the OFP was later in less favourable seasons, as later flowering allowed crops to escape frost. Higher yielding seasons also had a broader optimal period, and in the cases above largely overlapped with the optima in lower yielding seasons. In a practical sense, given seasonal conditions are unknown at sowing, wheat producers should aim to have crops flowering in the period bounded by 95% of the maximum yield achieved (Table 3).

**Figure 3:**
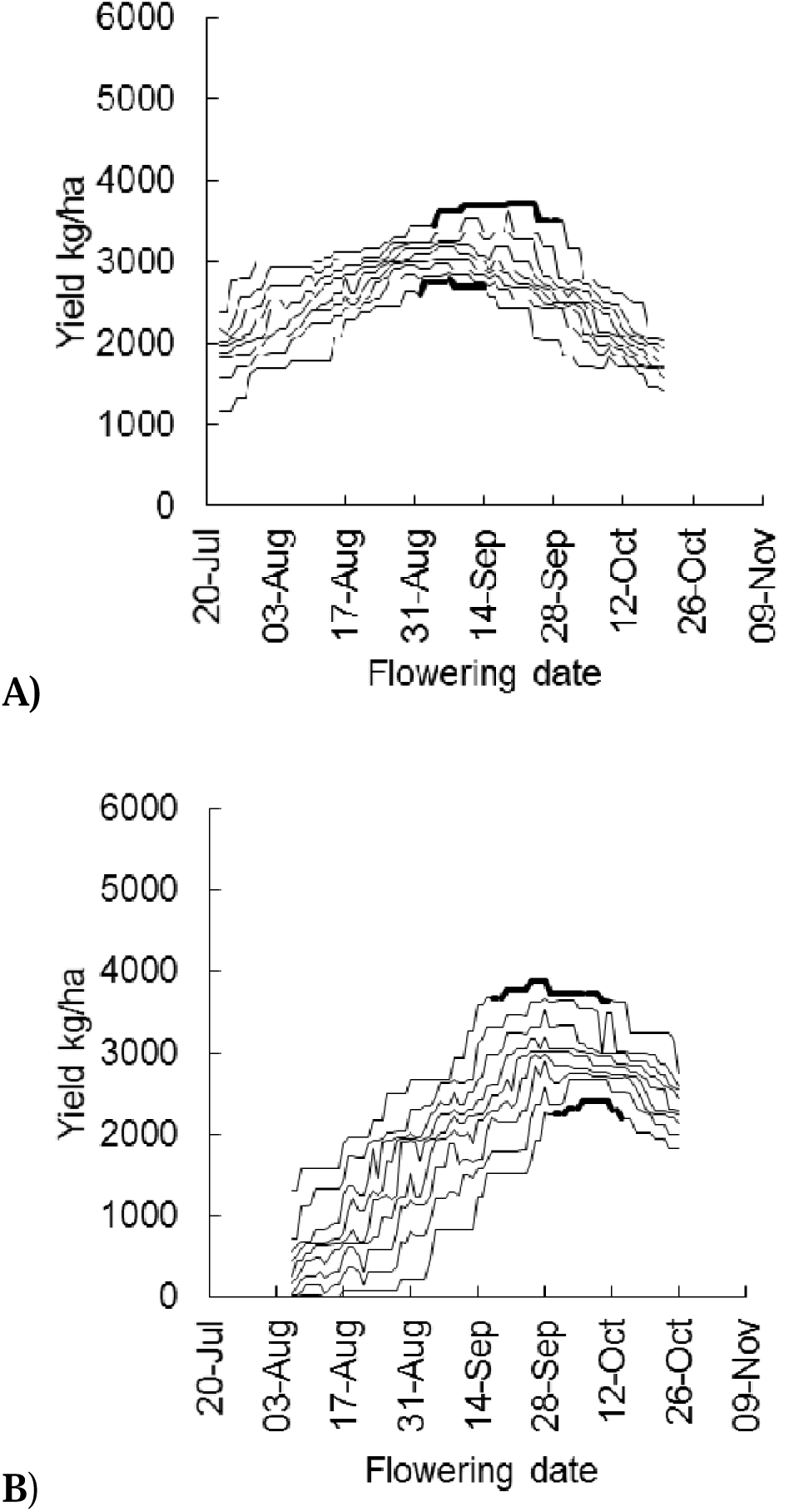
The relationship between frost and heat limited (FHL) yield (kg ha^-1^) and flowering date for a mid-fast cultivar of wheat at A) Lameroo, SA and B) Temora, NSW split into percentiles. Lines represent the simulated 10^th^, 20^th^, 30^th^, 40^th^, 50^th^, 60^th^, 70^th^, 80^th^ and 90^th^ percentiles of yield and corresponding flowering date values generated from 16 sowing dates over 51-years (1963-2013). Darkened line is the optimum flowering period for the 10^th^ and 90^th^ percentiles defined by ≥ 95% of the maximum mean yield.

Figure 4 shows how OFP have been defined using FHL yield. Figure 4 shows just 6 diverse locations, the same figures for all 28 locations can be found in Supplementary Figure 1. The degree of the incline or decline of the curves before or after the optimum for a location illustrates the influence of sub-optimal radiation and/or frost (before optimum), or accelerated development and/or drought and/or heat on grain yield (after optimum). For example, Minnipa (Fig. 4B) curves have a gentle incline showing that radiation and frost are less of a determinant on OFP, but the sharper decline after the optima shows heat and water stress play a larger role after peak yield is reached. In comparison, in locations such as Inverleigh and Dubbo (Fig. 4D and 4F), frost or sub-optimal radiation are greater determinants of the OFP, as seen by the sharp incline of the curves. The positive and negative standard deviation lines in Figure 4 show the variation around the mean, and reflect seasonal variability and the stability of a location’s environment.

**Figure 4:**
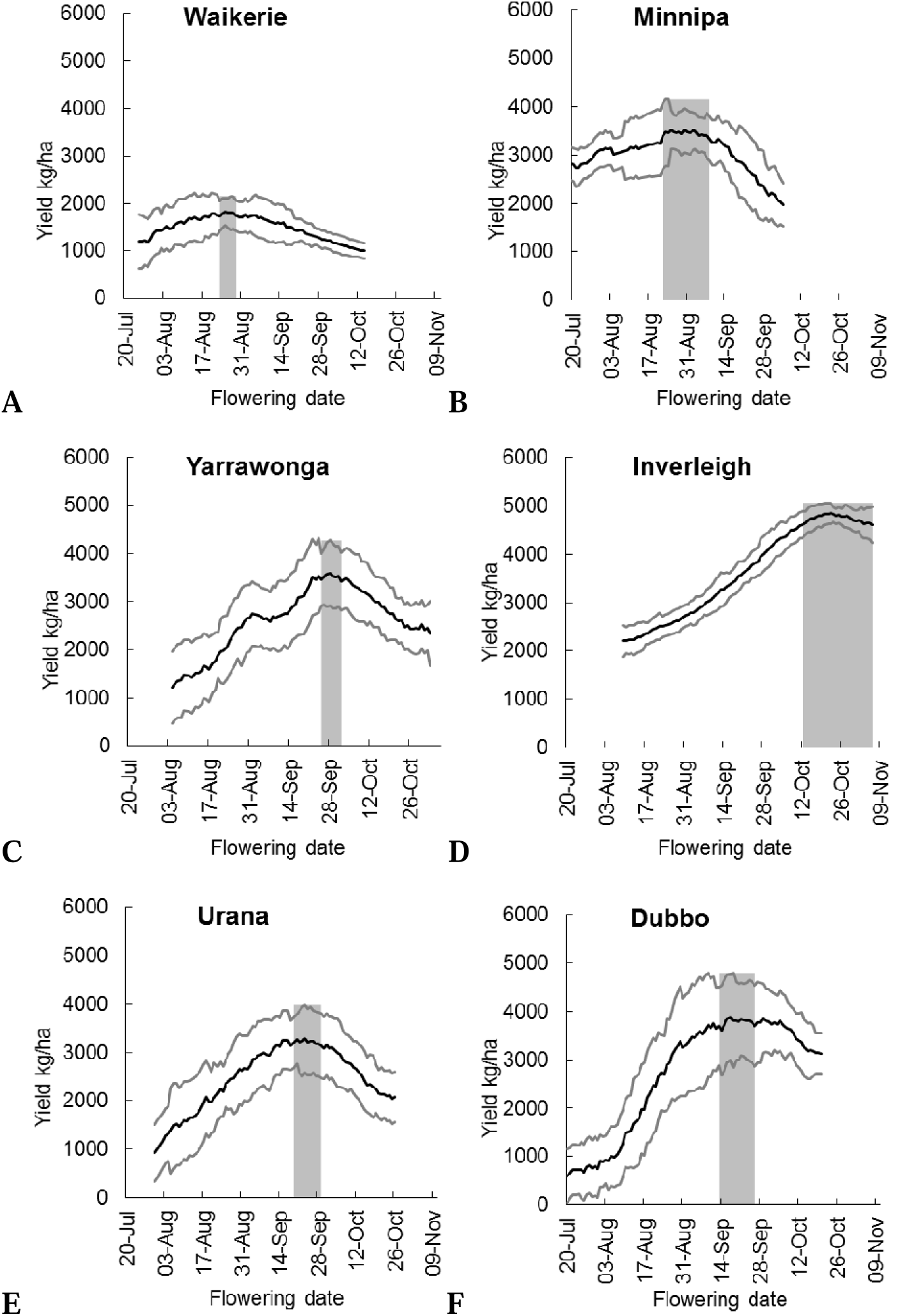
The optimal flowering period (OFP) for a mid-fast cultivar of wheat determined by APSIM simulation for A) Waikerie, SA B) Minnipa, SA C) Yarrawonga, VIC D) Inverleigh, VIC E) Urana, NSW F) Dubbo, NSW. Black lines represent the frost and heat limited (FHL) 15-day running mean yield (kg ha^-1^). Grey lines represent the standard deviation of the FHL mean yield (kg ha^-1^). Grey columns are the estimated OFP defined as ≥ 95% of the maximum mean yield achieved from the 51 seasons (1963-2013). All locations shown as in Figure 4 are located in Supplementary Figure 1.

In this analysis we compared two different soils at three locations; Swan Hill, Hopetoun and Bogan Gate (Table 1). In each instance, there was a heavier textured soil with higher plant available water capacity (PAWC), and a soil with a lower PAWC. At Swan Hill the OFP for the lighter soil (APSoil file 719) was 10 days shorter than for the heavier soil (APSoil file 718) (Table 3). In Hopetoun, the flowering period for the lighter soil began 3 days later but was of similar duration as that of the heavier soil (5 and 6 days respectively) (Table 3). The flowering period for the two soils at Bogan Gate were very similar (Table 3). There was some indication that OFP on low PAWC soils were earlier than for higher PAWC soils, presumably because higher PAWC soils were more able to buffer against terminal drought.

### 3.2 Relative importance of temperature extremes (frost and heat) and water stress in determining OFP

The OFP which has been defined for wheat in Figure 4 represents the combined effect of frost, heat and water stress and radiation on wheat physiological processes and yield. Figure 5 shows three of these sites again, but provides greater detail on how the OFP moves in response to these stresses, and how by using APSIM yield in addition to heat and frost rules gives a more accurate definition of the OFP, compared to using temperature extremes alone.

**Figure 5:**
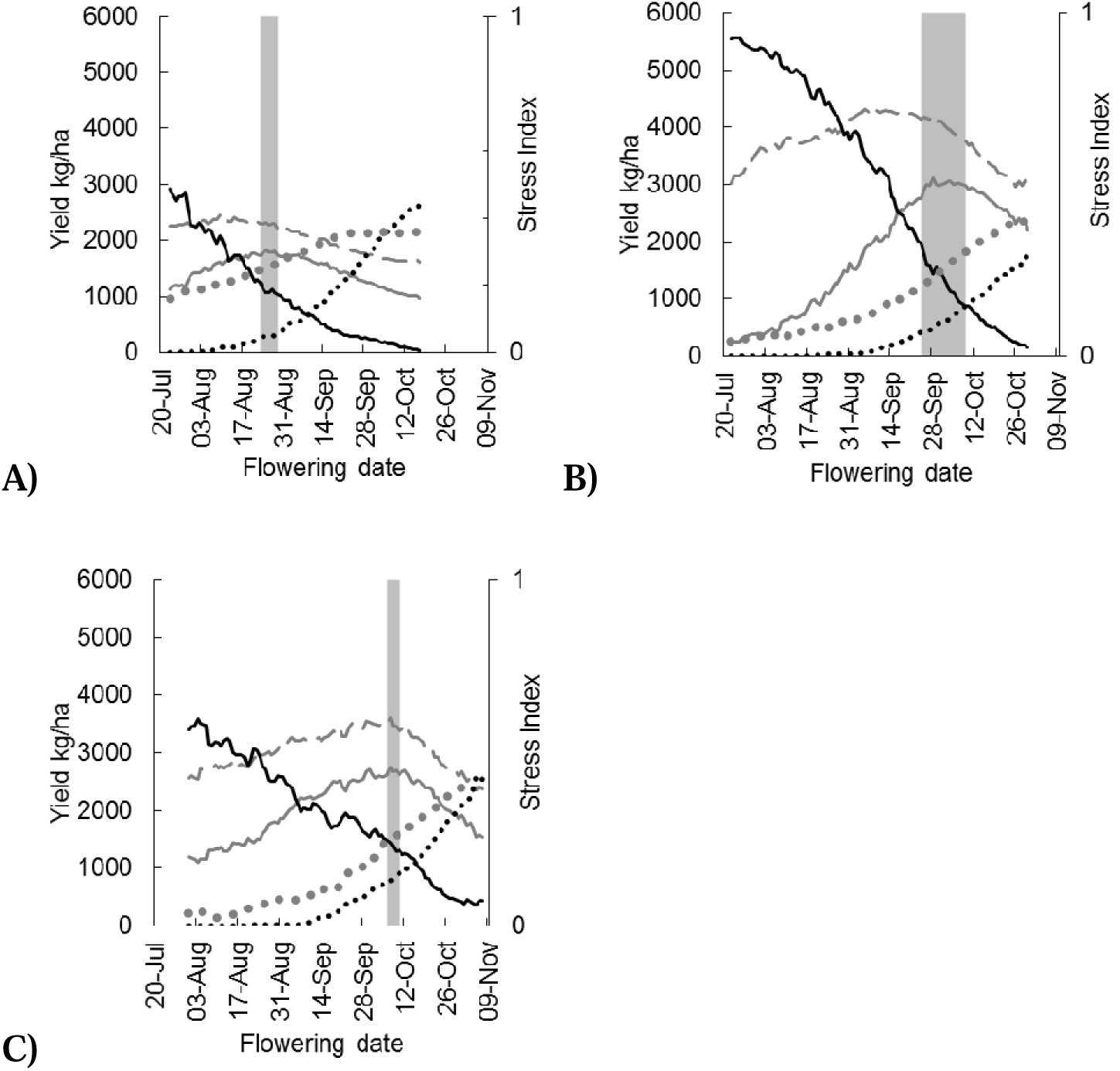
Potential yield (kg ha^-1^) (broken grey line), frost and heat limited yield (kg ha^-1^) (solid grey line) and the mean heat (dotted black line), frost (solid black line) and mean water stress from floral initiation to maturity (dotted grey line) indices (0 to 1) applied to yield plotted against flowering date for a mid-fast cultivar of wheat. A) Waikerie, SA B) Temora, NSW C) Longerenong, Vic. Grey columns are the estimated optimal flowering period defined as ≥ 95% of the maximum mean yield achieved in the 51 seasons (1963-2013).

Figure 5A for Waikerie demonstrates how using extreme temperatures alone to calculate OFP can result in severe overestimates in the timing of its occurrence. The point at which heat and frost damage lines intersect (where combined damage is lowest), is 8 September, however the flowering date for peak FHL yield is 25 August, 14 days earlier. The peak potential yield (APSIM without frost and heat damage) is earlier again, at 9 August. At Temora and Longerenong there is a similar pattern, although the shifts were less extreme (Fig. 5B, 5C). At Temora the intercept of the frost and heat indices is the 9 October, but the peak FHL yield is achieved from a flowering date on 28 September. Without applying the heat and frost indices, the peak potential yield was achieved by flowering on the 5 September. At Longerenong, the intercept of the frost and heat index is 15 October, but the peak FHL yield and peak potential yield is achieved from a flowering date on 7 October. In all of these Figures the effect of water stress during the reproductive phase in defining OFP is clear. Although only three examples are shown, similar patterns are observed at other sites included in this study. These examples clearly demonstrate that integration of both extreme temperatures and radiation, as well as water supply and demand as per our method are essential for accurate definition of OFPs.

### 3. 3 Implications for sowing time and genotype

Figure 6 and Table 4 illustrate that OFP for mid-fast, very-slow spring and winter cultivars are very similar in the environments where this comparison was made, but that the slower developing cultivars sown early have similar or higher yield potential.

**Figure 6:**
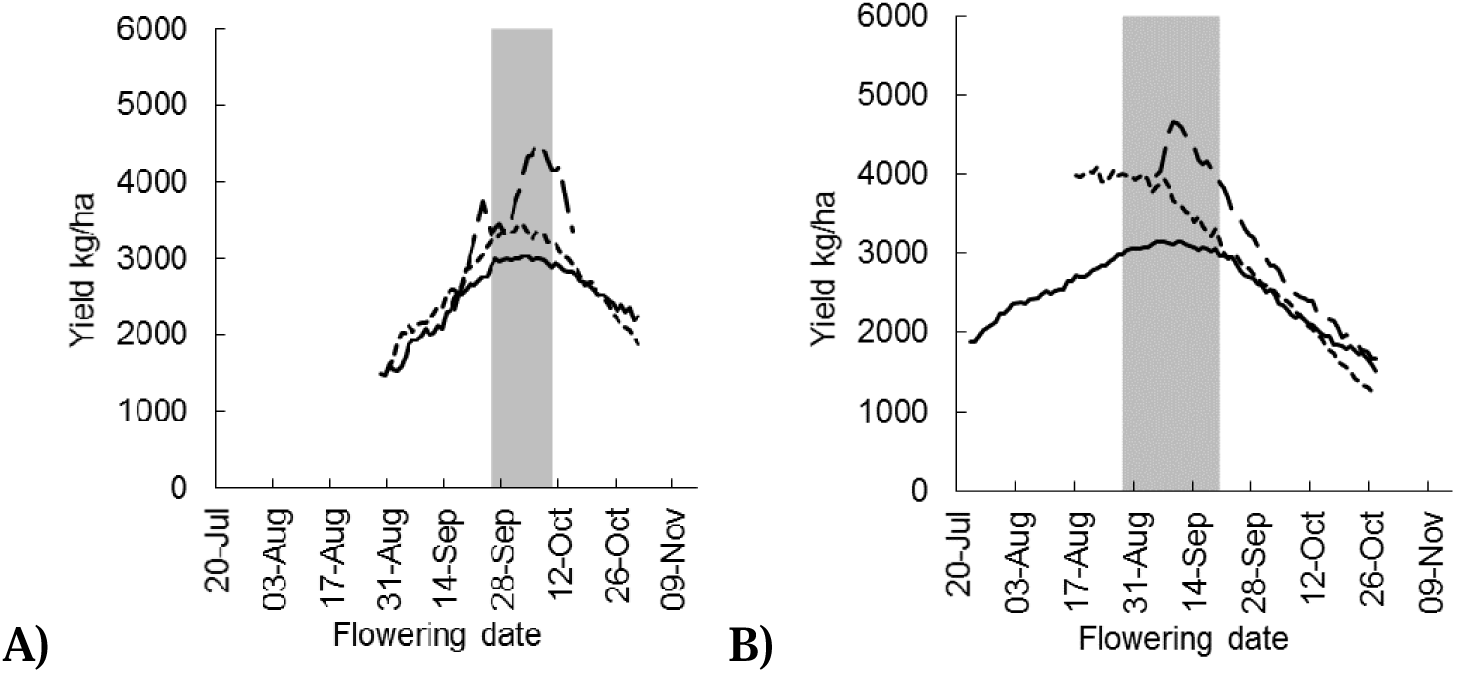
The 15-day running mean yield (kg ha^-1^) and flowering date of different wheat development types at A) Temora, NSW and B) Lameroo, SA over a 51-year simulation (1963-2013). Mean yields are for a mid-fast developing spring wheat (solid line), a very slow developing spring wheat (dotted line), and winter wheat (broken line). Grey columns are the estimated optimal flowering period defined as ≥ 95% of the maximum mean yield of a mid-fast cultivar.

**Table 4:** Simulated flowering date corresponding to peak mean frost and heat limited yield (kg ha^-1^) and sowing date for cultivars of different development type at Temora, NSW and Lameroo, SA.

| Maturity type | Temora |  |  | Lamerloo |  |  |
| --- | --- | --- | --- | --- | --- | --- |
|  | Peak mean yield (kg/ha) | Flowering date peak mean | Sowing date corresponding to peak yield | Peak mean yield (kg/ha) | Flowering date peak mean | Sowing date corresponding to peak yield |
| Winter | 4514 | 9-Oct | 12-Apr | 4653 | 9-Sep | 2-Apr |
| Very slow spring | 3444 | 2-Oct | 19-Apr | 4086 | 22-Aug | 22-Mar |
| Mid fast spring | 3038 | 3-Oct | 14-May | 3154 | 7-Sep | 6-May |

### 3. 4 Evidence of climate change

There is evidence that OFPs have shifted over the 51-year period considered in this study. At Lameroo the OFP for the years 1963-1997 was 4 to 19 September, Longerenong was 27 September to 2 October, and Charlton was 22 to 30 September. For the years 1998-2013 the OFP in Lameroo was 25 August to 8 September (Fig. 7A), Longerenong 8 to 11 September (Fig. 7B), and in Charlton 1-4 September (Fig. 7C). This can be attributed to lower than average rainfall and higher than average temperatures at flowering. Figure 8 illustrates climate change in Lameroo with a summary of September temperatures and growing season rainfall during the 51-year simulated time period.

**Figure 7:**
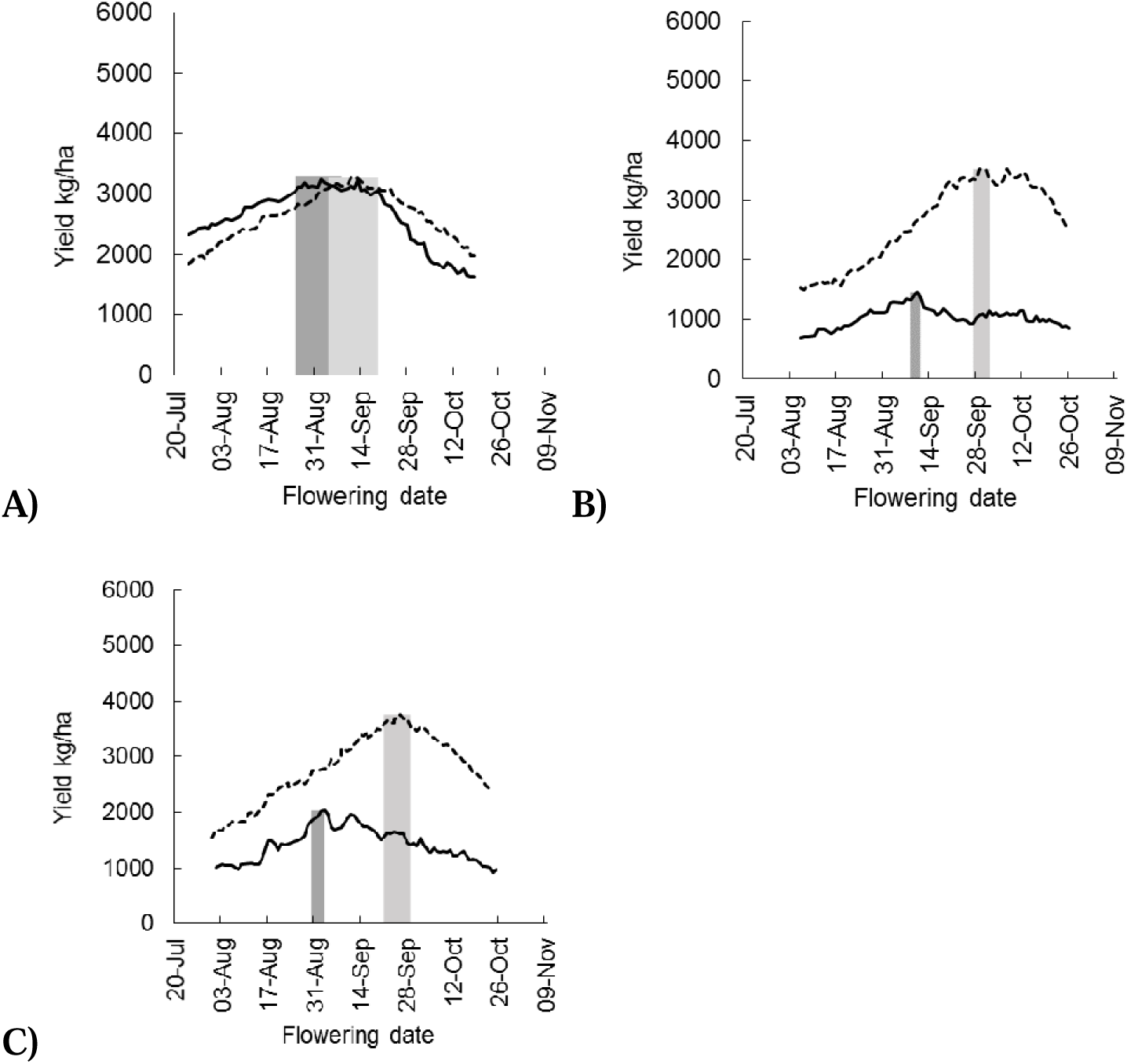
Estimated optimal flowering period (OFP) for mid-fast cultivar of wheat at A) Lameroo, SA and B) Longerenong, Vic C) Charlton, Vic for 35 years (1963-1997) (light grey column) and 16 years (dark grey column) (1998-2013). Mean frost and heat limited yield (kg ha^-1^) for 1963-1997 (broken line) and for 1998-2013 (solid line). Estimated OFP is defined by ≥ 95% of the maximum mean yield.

**Figure 8:**
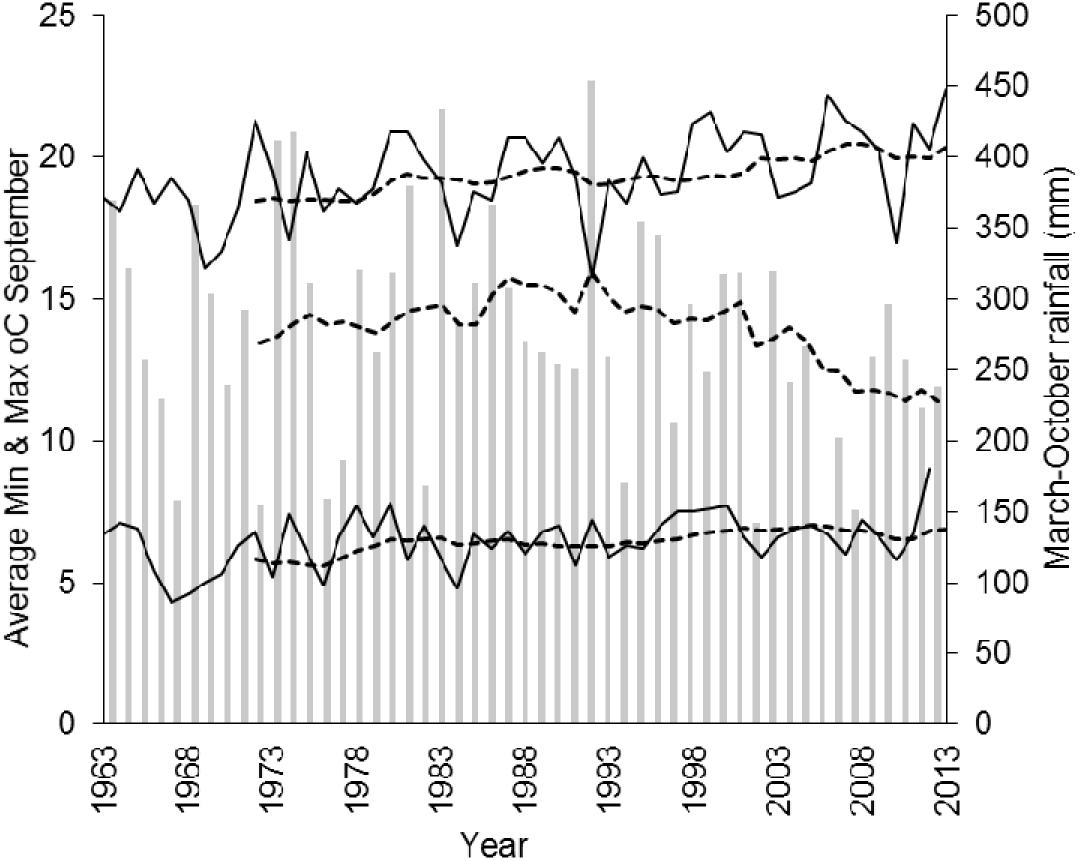
Growing season rainfall (columns) and average minimum and maximum temperature for September (solid lines) at Lameroo, SA. Broken lines are 10-year rolling means for rainfall and for minimum and maximum temperatures.

## 4. Discussion

### 4.1 Variation in optimal flowering periods

Variation in OFP duration indicated that defining OFP as 10 days either side of the optimum as proposed by Anderson *et al.* (1996) was not appropriate in all environments. Differences in the duration of OFPs were an indication of seasonal variability in a given environment, and the possible reasons causing the variability could be numerous. Wide OFPs may suggest that rainfall, frost and heat were more variable, as highest yields were achieved across a broader range of flowering dates e.g. Lameroo. Conversely wide OFP can also suggest a benign environment, resulting in a wide and stable environment for flowering e.g. Inverleigh. A narrower OFP suggests that an environment was less variable because optimal conditions for flowering occurred at approximately the same date in each season. Alternatively, a narrow period may indicate that the environment restricts the OFP due to overlap between the period of declining frost risk and the period of increasing drought and heat risk e.g. Hopetoun.

Zheng *et al.* (2012) also identified the OFP for two of the sites simulated here, Waikerie and Dubbo. At both sites the OFP in the Zheng *et al.* (2012) study were of a longer duration and later in the season than estimated here. While exact dates were not given in Zheng *et al.* (2012), for Waikerie the approximate OFP was the first week of September to mid-October, whereas by including water stress as a factor we found it to be much earlier and shorter from 23 August to 29 August. For Dubbo, Zheng *et al.* (2012) calculated the OFP from the last week of September to first week of November, while here it was 15 to 22 September. These differences in OFP reflect the methods used to calculate the OFP. Zheng *et al.* (2012) considered only the risk of heat and frost periods (i.e. temperature extremes) to define OFP, while here we also accounted for water stress and radiation effects on yield. This indicates that estimates of OFP defined only by extremes of temperature tend to be later in the season than when effects of drought are included.

In a two-year field study conducted by Gomez-Macpherson and Richards (1995), the optimal flowering date (determined by the highest yielding lines) for Condobolin was 10 September, and Penrose (1997) identified Temora’s optimum date as 1 October. In this study we found the OFP for Condobolin to be 11 to 19 September with peak yield falling on 15 September, and the OFP for Temora to be from 25 September to 10 October with peak yield falling on 3 October. In this comparison between experimental and our simulation method, the OFPs were very closely aligned, as in both cases optimal yield accounts for both temperature and water stress, rather than temperature extremes alone. However, our 51-year simulation study takes into account the long-term environmental patterns, providing a longer-term perspective.

### 4.2 Relative importance of temperature extremes (frost and heat) and water stress in determining OFP

Previous authors have defined OFPs by the occurrence of the last frost event and first heat event (Zheng *et al.*, 2012). We have shown that OFPs are actually defined by a trade-off between drought, radiation, frost and heat, rather than temperature as the primary factor (Fig. 5). Any crop flowering on the optimal date for yield in a given combination of site and season may experience all three abiotic stresses to varying degrees during their critical period. The OFP we have presented represents the period that minimises the combined yield reduction from all three stresses. To our knowledge, this is the first comprehensive study to identify OFP for wheat across south-eastern Australia that captures the impact of these stresses across a wide range of climate and soil-type variation. The approach would be applicable to other crops (e.g. barley, canola, grain legumes) where yield is also dependant on flowering in an optimal period.

### 4.3 Implications for sowing time and genotype

Defining OFP is useful because it allows identification of appropriate sowing date x genotype combinations to optimise yield for a specific environment (Anderson *et al.*, 1996). Timing of flowering in a given environment is a function of sowing date, genotype (cultivar) and prevailing seasonal temperatures. Sowing date and cultivar selection are two major management decisions made by wheat producers at the start of the growing season, the objective being to select combinations that lead to crops flowering during the OFP. Throughout the history of wheat cultivation, farmers and wheat breeders have selected developmental patterns which best match the conditions under which they are grown (Cockram *et al.*, 2007; Kamran *et al.*, 2014). Since the introduction of photoperiod insensitive semi-dwarf wheat cultivars during the 1970s, breeding programs in SEA have focussed overwhelmingly on spring wheats of mid-to fast-maturity (Davidson *et al.*, 1985; Eagles *et al.*, 2009). The development pattern of these cultivars matched the timing of once reliable autumn planting rains in April-May with OFPs in spring. However, optimal sowing times for mid-fast cultivars (predominantly April-May, Table 3) which allow them to flower in the OFPs unfortunately coincides with the most severe period of recent rainfall decline identified by Cai *et al.* (2012) and Pook *et al.* (2009). New combinations of genotype x management will be required to ensure crops continue flowering in the OFPs. Strategies could include using stored soil water from either summer rainfall (Hunt *et al.*, 2013; Hunt and Kirkegaard, 2011; Verburg *et al*., 2012), or long fallow (Oliver *et al.*, 2010) to establish crops. This practice has been shown to be successful in low rainfall locations in the Pacific Northwest of the USA (Schillinger and Young, 2014) and has previously been proposed for SEA by Kirkegaard and Hunt (2010). This practice would be facilitated by using genotypes with long coleoptiles which would allow them to emerge on stored soil water from greater soil depths (Rebetzke *et al.*, 2007). Another option would be to use winter wheat genotypes which can be established much earlier than spring genotypes when soil water is available, but still flower during the OFP as demonstrated by Penrose and Martin (1997). Establishment would then occur prior to the traditional sowing window, as late summer and early autumn rainfall has been more reliable in recent decades (Hunt and Kirkegaard, 2011). Previous simulation and field experiments have demonstrated that winter genotypes sown early yield as well as (Frischke *et al.*, 2015; Gomez-Macpherson and Richards, 1995; Kirkegaard *et al.*, 2014) or better (Bell *et al.*, 2015; Coventry *et al.*, 1993; Moore, 2009) than spring genotypes sown later. Figure 6 and Table 4 support these findings. Now that OFP have been accurately defined, breeders and agronomist can use our definitions to target OFP in any given season, and depending how the season breaks, provide growers with new genotype x sowing strategies, such as listed above, for specific environments to maximise yield in a variable climate.

The sowing date ranges targeting OFP estimated in this study (Table 3) are consistently of shorter duration than those currently recommended by state departments of agriculture. For example, the recommended sowing date of mid-fast spring maturity types sown in the Victorian Mallee is between the last week of April, and the second week of June (Department of Economic Development Jobs Transport and Resources and GRDC, 2015). According to our simulations, Kerang, situated in the Victorian Mallee has an optimal sowing date range of 25 April to 9 May, with a median sowing date of May 3. Similar conclusions can be reached at other sites in this analysis such as Longerenong in the Wimmera region of Victoria, Charlton in North Central Victoria and Yarrawonga in North Eastern Victoria, and also some sites in NSW (Matthews *et al.*, 2015). Whilst state recommendations appear to correctly estimate the start of the sowing period, they greatly overestimate the end of the optimal period. Currently no “sowing date guide” exists for South Australian farmers, but we believe trends to be similar there.

### 4.4 Climate change impacts on flowering and sowing dates

Global minimum and maximum temperatures are increasing (IPCC, 2013). By 2030, Australian temperatures are predicted to rise between 1.9^o^C and 5^o^C (CSIRO and Bureau of Meteorology, 2015). Figure 7 shows that OFPs have shifted at three sites, Lameroo, Longerenong and Charlton over the 51-year period considered in this study. A shift to an earlier OFP can be expected in other areas of the SEA wheat-belt where autumn rainfall decline and increased spring drought has been experienced in the period 1998-2013. APSIM has been used to analyse crop development under future climates in numerous studies. Zheng *et al.* (2012) studied wheat development under future climate scenarios, and estimated that both sowing and flowering dates would occur earlier. Yang *et al.* (2014) also predicted through simulation that flowering will be 11 days earlier on average in the Australian wheat belt. This was explained by earlier last frost days and earlier first heat events in future climates. However, predictions of increased minimum temperatures and decline in frost incidence for SEA incorporated in the studies of Zheng *et al.* (2012) and Yang *et al.* (2014) are at odds with observed increases in the occurrence and severity of late-season frosts across the region (Crimp *et al.*, 2015). The Millennium drought (1996-2009) occurred within the 1998-2013 simulated time period, and was characterised by lower than average rainfall and higher temperatures (Fig. 8). Our study has demonstrated that in low rainfall sites and seasons, highest yields were achieved when the wheat crops flowered earlier, escaping spring drought (Fig. 2, 3, 5 and 7). This emphasizes the importance of considering patterns of water supply and demand when defining flowering periods rather than temperature extremes alone.

## 5. Conclusion

OFPs for wheat in SEA vary with site and season and are largely driven by seasonal water supply and demand, with extremes of heat and temperature having a secondary although auto-correlated effect. Sowing dates required for current cultivars to achieve OFPs coincide with a period of marked autumn rainfall decline which has reduced grain yields across the SEA region. Further work to identify new G x M strategies that target OFPs which account for water and temperature stress is required, to avoid current and future yield losses associated with autumn rainfall decline.

## Supplementary Data

**Figure S1:** Optimal flowering period of wheat for 22 locations simulated and not displayed in Figure 4.

## Acknowledgements

This research was funded by GRDC through the Grains Industry Research Scholarship and project number CSP00174.

